# Vγ9Vδ2 T cells are potent inhibitors of SARS-CoV-2 replication and exert effector phenotypes in COVID-19 patients

**DOI:** 10.1101/2022.04.15.487518

**Authors:** Laetitia Gay, Marie-Sarah Rouviere, Soraya Mezouar, Manon Richaud, Laurent Gorvel, Etienne Foucher, Loui Madakamutil, Bernard La Scola, Amélie Menard, Jérôme Allardet-Servent, Philippe Halfon, Paul Frohna, Carla Cano, Jean-Louis Mege, Daniel Olive

## Abstract

Vγ9Vδ2 T cells play a key role in the innate immune response to viral infections, including SARS-CoV-1 and 2, and are activated through butyrophilin (BTN)-3A. Here, the objectives were to: 1) characterize the effects of SARS-CoV-2 infection on the number, phenotype, and activation of Vγ9Vδ2 T cells in infected patients, and 2) assess the effects of *in vitro* SARS-CoV-2 infection on the expression of BTN3A and its impact on the activation and response of Vγ9Vδ2 T cells to an anti-BTN3A antibody. Blood Vγ9Vδ2 T cells decreased in clinically mild SARS-CoV-2 infections compared to healthy volunteers (HV). This decrease was maintained up to 28 days and in the recovery period. Terminally differentiated Vγ9Vδ2 T cells tend to be enriched on the day of diagnosis, 28 days after and during the recovery period compared to HV. Furthermore, these cells showed cytotoxic and inflammatory activities as shown by TNFα, IFNγ and CD107a/b increase following anti-BTN3A activation. Moreover, BTN3A upregulation and Vγ9Vδ2 T cell infiltration were observed in a lung biopsy from a fatal SARS-CoV-2 infection, as compared to HV. *In vitro*, SARS-CoV-2 infection significantly increased BTN3A expression in macrophages and lung cell lines. The activation via BTN3A enhanced the anti-SARS-CoV-2 Vγ9Vδ2 T cells cytotoxicity and IFN-γ and TNFα in SARS-CoV-2 infected patient. Increasing concentrations of anti-BTN3A were accompanied by an inhibition of viral replication. Altogether, these data suggest that Vγ9Vδ2 T cells are important in the immune response against SARS-CoV-2 infection and that activation by an anti-BTN3A antibody may enhance their response.

**KEY POINTS:**

- SARS-CoV-2 mediates upregulation of the key receptor of Vγ9Vδ2 T cells BTN3A on lung tissues and cell lines as well as monocytes
- During SARS-CoV-2 infection, Vγ9Vδ2 are differentiated and efficiently degranulate and secrete cytokines upon activation with BTN3A mAb

## INTRODUCTION

The global pandemic caused by the emerging severe acute respiratory syndrome coronavirus-2 (SARS-CoV-2) has resulted to date in more than 456 million global cases of coronavirus disease (COVID)-19 with more than 6 million deaths^1^. The disease is characterized by very heterogeneous clinical manifestations. While some infected persons remain asymptomatic, some will develop a mild respiratory disease that will resolve with no or little medical attention, and others (10 to 20% of symptomatic persons), will experience a severe disease often occurring as a sudden deterioration ~11 days after the onset of first symptoms^2^, associated with respiratory failure and multi-organ complications possibly leading to patients’ deaths^3,4^. A post-COVID syndrome with unrelenting symptoms seems to occur in 10 to 15% of patient^5^. Unbalanced immune responses have been linked to the disease course and severity, and may be involved in chronic COVID-19 symptoms^4^. Early publications described lymphopenia mostly affecting T and B cells, neutrophilia and decreased eosinophil and monocyte counts^6,7^. The percentage of Vγ9Vδ2 T cells in COVID-19 patients at the time of hospital admission has been reported to be significantly lower than that of matched healthy volunteers (HV), with the relationship being strongest in patients with a fatal outcome^8,9^. This suggests the involvement of the Vγ9Vδ2 T cells in the in COVID-19 disease and their role needs further characterization.

Vγ9Vδ2 T cells, known for their potent elimination of cells with intracellular bacterial infection, have also strong antiviral activities^10^. Two direct antiviral actions of Vγ9Vδ2 T cells include cytotoxic activity to pathogen-infected cells and a cell-mediated non-cytolytic activity based on cytokine production and release^10–12^. During the SARS-CoV-1 2003 outbreak, Vγ9Vδ2 T cells were shown to inhibit SARS-CoV-1 replication *in vitro* through an IFNγ-dependent process^13^. In SARS-CoV-1 infected patients, the effector memory Vγ9Vδ2 T cell population was selectively expanded in the peripheral blood 3 months after the onset of the disease^13^ possibly contributing to adaptive responses.

In humans, Vγ9Vδ2 T cell responses are triggered by the cell surface molecule butyrophilin-3A1 (BTN3A1) that will recognize through its intracellular domain phosphorylated nonpeptide antigens, called phosphoantigens (pAgs), which are overproduced intracellularly in infected or tumor cells or naturally produced by several pathogens^14,15^.

In this study, our aim was to characterize the effects of SARS-CoV-2 infection on the number, phenotype, activation status and responses of Vγ9Vδ2 T cells from COVID-19-patients during the active infection period and during recovery. Next, we characterized the infiltration of Vγ9Vδ2 T cells and the expression of BTN3A in the lungs of patients. Finally, we quantified the effects of SARS-CoV-2 infection on the expression of BTN3A in infected cells and characterized its impact on the activation and responses of Vγ9Vδ2 T cells *in vitro*.

## METHODS

### Study participants and ethics statement

This prospective observational study (NCT04816760) was conducted in accordance with the Declaration of Helsinki and the French law on research involving humans (ANSM, ID-RCB: 2020-A00756-33). The protocol of the study was approved by an independent national review board (Comité de Protection des Personnes, Ile-de-France XI, ID: 20027-60604) and registered at ClinicaTrials.gov (NCT04816760). All patients provided written informed consent. Patients were recruited at the European Hospital and the North Hospital of Marseille (France) from March 30 to December 20, 2020 and from November 2020 to June 2021 for the first and second cohort, respectively. Healthy volunteers (HV)’s samples were obtained from the Etablissement Français du Sang (EFS Alpes-Méditerranée).

### Clinical samples

COVID-19 patients in the acute phase (Kinetics patients): Ten patients with confirmed SARS-CoV-2 infection (positive RT-PCR on nasopharyngeal swab) were included within the first 3 days of their hospital admission either during the first wave (n=6 from April 07 to April 20, 2020) or the second wave (n=4 from November 14, 2020 to April 18, 2021) of the pandemic. All patients had lung computed tomography scans showing evidence of COVID-19 pneumonia on the day of admission. The clinical severity of the pneumonia was assessed using the WHO progression scale^16^. All patients were managed within the COVID-19 ward except one patient which required intensive care unit admission during his stay. Biological samples were collected at 4 time points over the first month after inclusion: Day 0, 7, 14 and 28 (D0, D7, D14 and D28). Among patients of the second wave, an additional time point at D90 was obtained.

COVID-19 patients in the post-COVID (PC) infection phase: Seven patients were investigated after a symptomatic SARS-CoV-2 infection, during the recovery phase which ranged from D59 to 92.

### Cell Isolation and cells lines cultures

Peripheral blood mononuclear cells (PBMCs) were isolated from patient’s peripheral blood or from 15 HV buffy coats by density-gradient centrifugation using Ficoll (Eurobio) at 800 × *g* (brake off) for 30 min at 20°C. PBMCs were cryopreserved at −80°C in 90% fetal bovine serum (FBS, Gibco) + 10% dimethyl sulfoxide (DMSO, Sigma-Aldrich).

Monocytes were purified from PBMCs through a CD14 selection using MACS magnetic beads (Miltenyi Biotec) and cultured in Roswell Park Memorial Institute-1640 medium (RPMI, Life Technologies) containing 10% FBS, 2 mM L-glutamine, 100 U/mL penicillin and 50 μg/mL streptomycin (Life Technologies). For macrophages-derived from monocytes (MDMs), cells were cultured in RPMI-1640 containing 10% inactivated human AB-serum (MP Biomedicals), 2 mM glutamine, 100 U/mL penicillin and 50 μg/mL streptomycin during 3 days. Then, the medium was replaced by RPMI-1640 containing 10% FBS and 2 mM glutamine, and cells were differentiated into macrophages for 4 additional days.

Vγ9Vδ2 were expanded from fresh PBMCs. Briefly, PBMCs were grown in RPMI-1640 supplemented with 10% FBS, interleukin-2 (IL-2, 200 UI/ml) and Zoledronic acid monohydrate (final concentration of 1 μM). IL-2 was added every 2 days beginning on day 5. After being cultured for 12 days, the purity of the Vγ9Vδ2 was assessed by flow cytometry analysis (> 85%) and then frozen at −80°C in 90% FBS + 10% DMSO.

Human bronchial epithelial cells (BEAS-2B cells, ATCC® CRL-9609™) were cultured in LHC-9 medium (Life Technologies) and human lung fibroblast (MRC-5 cells, ATCC® CCL-171™) were cultured in Minimum Essential Media (MEM, Life Technologies) supplemented with 4% FBS and 2 mM L-glutamine at 37°C in an atmosphere of 5% CO_2_.

### Virus production and cell infection

SARS-CoV-2 strain IHU-MI6 was obtained after VeroE6 cells (ATCC® CRL-1586™) infection in MEM supplemented with 4% FBS as previously described^17^. Monocytes, MDMs, BEAS-2B and MRC-5 cells were infected with virus suspension at multiplicity of infection (MOI) of 1 for 24 hours at 37°C in an atmosphere of 5% CO_2_.

### Immune population characterization

This was performed on the first cohort patients. Cryopreserved PBMCs were thawed, washed and incubated at 37°C for 30 min. Rested PBMCs were filtered using 30 μm filter, washed and a minimum of 200,000 cells were seeded into wells of a 96-well plate (U bottom). Cells were then stained for spectral flow cytometry (**Supplemental Table 1**).

### Vδ2 T cell stimulation and functional assessment

This was performed on the second cohort patients. Cryopreserved PBMCs were thawed, washed and rested overnight in cell culture supplemented or not with 100U/ml IL-2 (Proleukin®, Clinigen). Rested PBMCs were filtered using 30 μm filter, washed and a minimum of 200,000 cells were seeded into wells of a 96-well plate (U bottom). PBMCs cultures not supplemented with IL-2 overnight were stimulated with 25 ng/ml phorbol 12-myristate-13-acetate (PMA) (Sigma-Aldrich) and 1 μg/ml ionomycin (Sigma-Aldrich). Subsequently, PBMCs which were supplemented with IL-2 overnight were stimulated with 5 μg/ml anti-BTN3 20.1 mAb (Imcheck Therapeutics). Unstimulated PBMCs control was also added with medium only. All conditions were incubated in the presence of 0.3 μl/well brefeldin A (GolgiPlug, BD Biosciences) and Phycoerythrine (PE)-conjugated anti-CD107a and anti-CD107b. Plates were briefly centrifuged and incubated for 4 hours at 37°C. Cells were then stained for spectral flow cytometry (**Supplemental Table 2**).

### Flow Cytometry staining and data acquisition

PBMCs were stained with antibodies as stated in **Supplemental Table 1 or Table 2** for the first (phenotype) or second cohort (function), respectively. Extracellular staining procedure was performed at room temperature. PBMCs were washed twice and incubated 15 min with viability dye (LIVE/Dead^TM^ Fixable Blue Viability kit, Invitrogen). Cells were washed twice and incubated first in presence of Fc Block (BD Biosciences) and CC-chemokine receptor 7 for 10 min, and then with other monoclonal antibodies diluted in Brilliant Stain Buffer Plus (BD Biosciences) for 20 min. Samples were washed and the pellet suspended in 150 μl phosphate-buffered saline (PBS) with 1% Cytofix (BD Biosciences) for the first cohort. For the second cohort, fixation/permeabilization kit (BD Biosciences) was used for intracellular staining with Allophycocyanin (APC)-TNFα and Fluorescein Isothiocyanate (FITC)-IFNγ. Samples were washed, and the pellet suspended in 150 μl PBS. Samples were then acquired with a maximum available event on a Cytek®Aurora cytometer (Cytek Biosciences) with the SpectroFlo®software (Cytek Biosciences).

For *in vitro* assays, SARS-CoV-2 infected-cells were suspended in PBS (Life Technologies) containing 1% FBS and 2mM EDTA (Sigma-Aldrich). Cells were labeled with viability dye (LIVE/Dead™ Fixable Near-IR Stain Kit) and with anti-BTN3A (103.2) or the isotype control (Miltenyi Biotech). After 30 min incubation, primary antibodies binding were detected with Alexa Fluor-488 anti-mouse (Invitrogen). Data were collected on a BD Canto II instrument (BD Biosciences) and analyzed with FlowJo software (FlowJo v10.6.2, Beckton Dickinson).

### Flow cytometry data analysis

Data cleaning was performed on SpectroFlo®software (Cytek Biosciences) by excluding doublets and dead cells on the CD45^+^ leukocyte population. Next, lymphocytes were analyzed with OMIQ software using the following pipeline: (1) verifying and modifying scaling of the data to ensure the hyperbolic arcsine (arcsinh) transformation was unimodal around 0, (2) manual gating to identify the main populations and (3) subsampled at 9,000 cells for the first cohort and 1,000 cells for the second cohort before UMAP (Uniform Manifold Approximation and Projection) running (neighbors: 15, minimum distance: 0.4, components: 2, metric: euclidean). FlowJo software was used for manual gating and statistical analysis.

### Immunohistofluorescence

A lung biopsy sample was obtained from a mechanically ventilated patient with severe COVID-19 pneumonia after 34 days of evolution. The lung sample was free from SARS-CoV-2 mRNA and was embedded in optimal cutting temperature (OCT) before freezing. Healthy lung tissue collected before the pandemic and tonsil tissue were also embedded in OCT. The 8-10 μm tissue sections were cut with cryostat (NX70 MicromMicrotech) and we used the Tyramide SuperBoost™ kit (Invitrogen) according to the manufacturer’s instructions for immunofluorescence staining. 5 μg/ml IgG1 mouse monoclonal antibody 20.1 directed against BTN3A (Imcheck Therapeutics) or 10 μg/ml IgG1 mouse monoclonal antibody BB3 directed against Vδ2-TCR (kind gift of Dr. Pende, Genoa, Italy supplier) were used as primary antibodies. 100 μl of the antibody dilution per slide was sufficient. Alexa Fluor™ Tyramide, which reacts with HRP, allowed signal amplification for the detection of these low-abundance targets. A second staining with 10 μg/ml IgG2b mouse monoclonal antibody 9C4 directed against EpCam (Biolegend) allowed the identification of epithelial cells by indirect detection with anti-IgG2b Cyanine-3 labelled secondary Ab (Jackson ImmunoScience, dilution 1:500). The sections were washed in PBS and incubated 10 min with 4’,6-diamidino-2-phenylindole (DAPI, 1/10,000, Sigma-Aldrich) to visualize the cell nuclei. The images were acquired on confocal microscope (Zeiss LSM 880 airy scan) using Zen Black software (Zeiss) and Fiji software (ImageJ-win64) used for data treatment.

### RNA isolation and q-RTPCR

Total RNA was extracted from monocytes, MDMs, BEAS-2B and MRC-5 (2×10^6^ cells/well) using the RNeasy Mini Kit (Qiagen) with DNase I treatment to eliminate DNA contaminants as previously described^18^. The quality and quantity of the extracted RNAs were evaluated using a spectrophotometer (Nanodrop Technologies). Reverse-transcription of isolated RNA was performed using a Moloney murine leukemia virus-reverse transcriptase kit (Life Technologies) and oligo (dT) primers. The expression of *BTN3A* isoform genes was evaluated using real time qPCR, Smart SYBR Green fast Master kit (Roche Diagnostics) and specific primers (**Supplemental Table 3**). The qPCRs were performed using a CFX Touch Real-Time PCR Detection System (Bio-Rad). The results were normalized by the expression of *ACTB* housekeeping gene and are expressed as relative expression of investigated genes with ΔCt = Ct_Target_ – Ct_*ACTB*_ as previously described^19^.

### Viral RNA extraction and q-RTPCR

Viral RNA was extracted from infected monocytes, MDMs, BEAS-2B and MRC-5 using NucleoSpin® Viral RNA Isolation kit (Macherey-Nagel). One-Step RT-PCR SuperScript™ III Platinum™ Kit (Life Technologies) was used for virus detection. Thermal cycling was achieved at 55°C for 10 min for reverse transcription, pursued by 95°C for 3 min and then 45 cycles at 95°C for 15 sec and 58°C for 30 seconds using a LightCycler-480 Real-Time PCR system (Roche). The primers and the probes were designed against the E gene^17^.

### Cytotoxicity assay

Monocytes, MDMs, BEAS-2B and MRC-5 were labeled with 10 μM Cell Proliferation Dye eFluor® 670 (Invitrogen) and then stimulated with virus. Target cells were co-cultured with Vγ9Vδ2 T cells (effector) at 1:1 effector-to-target (E:T) ratio in presence of anti-BTN3A 20.1 Ab (0, 0.1, 1 or 10 μg/ml). After 24 hours, cells were stained with CellEvent Caspase-3/7 Green (Invitrogen). The cytotoxicity was assessed by flow cytometry as the percentage of Caspase 3/7^+^ cells in the target cell population. Data were collected on a BD Canto II instrument (BD Biosciences) and analyzed with FlowJo software.

### Immunoassays

Monocytes, MDMs, BEAS-2B and MRC-5 were stimulated with virus and co-cultured with Vγ9Vδ2 T cells (effector) at 1:1 E:T ratio in the presence of anti-BTN3A 20.1 Ab (0, 0.1, 1 or 10 μg/ml). After 24 hours, culture supernatants from co-cultures were collected, and IFNγ and TNFα levels were measured with appropriate immunoassay Kits (R&D Systems) according to the manufacturer’s instructions.

### Statistical Analyses

Statistical analysis was performed using GraphPad Prism (GraphPad Software), for transcriptional analysis using the Student t or nonparametric Mann-Whitney U test and for Spectral Cytometry using non parametric Kruskal-Wallis test, followed by Dunn’s multiple comparisons test or two-way ANOVA, followed by Holm-Sidak’s multiple comparisons test. The limit of significance was set as *p*<0.05.

## RESULTS

### Long-lasting major decrease of Vγ9Vδ2 T cells in patients with COVID-19

The cellular kinetics (D0 to D28) of the first cohort of 6 patients in acute infection and 7 samples post-COVID-19 (PC) patients were included during the first wave of the pandemic. Baseline characteristics of the patients are presented in **Table 1**. We first investigated the dynamics of the different immune cell subsets using multicolor spectral cytometry panel (**Supplemental Table 1**), which identifies CD4, CD8, Vγ9Vδ2 and regulatory T cells, B cells, NK cells and monocytes. The dimensionality reduction by UMAP analysis (**Figure 1A**) showed a distribution of the main lymphocyte subsets according to marker co-expressions. Through this analysis as well as a manual gating (**Supplemental Figure 1**), we identified a reconfiguration of peripheral immune cell frequency. Specifically, a significant decrease of Vγ9Vδ2 T cells in the blood of patients with mild infection, as compared to healthy volunteers (HV), was observed at D0: 3.2 *vs*. 50.1%, *p*=0.0057 and D7: 4.5%*vs*. 50.1%, *p*=0.0199 (**Figure 1B**), while the decreases at D14 and D28 did not reach significance. In contrast, no significant difference from HV were observed in the frequency of CD8 conventional T cells (T conv) and the total γδ T cell population in COVID-19 patients (**Supplemental Figure 2A**).

**Figure 1.**
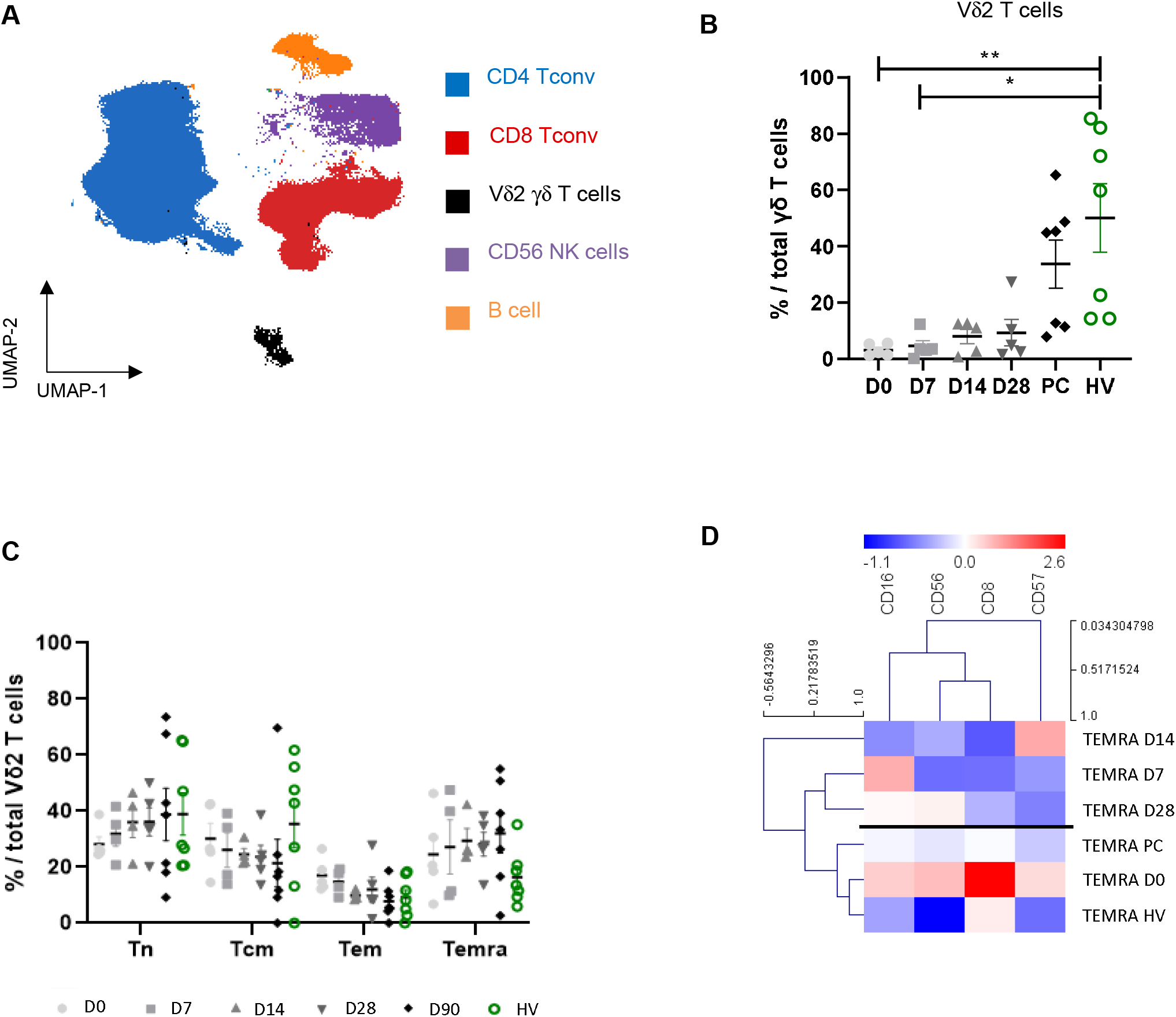
Long-lasting major decrease of Vγ9Vδ2 T cells in patients with COVID-19. (**A**) Manual gating clusters projected on to UMAP in OMIQ software. Subsampling with 50, 000 cells of post-COVID (PC) consensus file (n=5) was illustrated. The following markers were used to identify the immune cell populations: CD19 and CD20 for B cells (orange), CD16 and CD56 for NK cells (purple), CD3, CD4 (blue) and CD8 (red) for the main T cell populations, Vδ2 TCR for Vδ2 T cells (black). (**B**) Summary statistical graphs showing the alteration of Vδ2 γδ T cells frequencies. Dot plot include n=5 COVID-19 patients, n=7 PC patients and n=7 healthy volunteers (HV). Data were analyzed using Kruskal-Wallis test, Dunn’s multiple comparisons test; \**p*<0.05 and \*\**p*<0.01. (**C**) Dot plot of maturation Vδ2 γδ T cell subpopulations, identified by manual gating using CD27 and CD45RA markers. Results were showed for the patients (gray-scale kinetics) and HV (green). For Vδ2 γδ T cells maturation, kinetics of two patients at D7 was removed because Vδ2 γ T cells were less than 30 cells for analysis. Data were analyzed using two-way ANOVA and Holm-Sidak’s multiple comparisons test. (**D**) Shown representative heatmap of Temra Vγ9Vδ2 T cells maturation. Blue indicates low cellular frequency and red indicates high cellular frequency. The dendrograms for markers (columns) and samples (rows) were constructed with hierarchical clustering (Pearson’s correlation). Normalized expressions of each marker are from averages by kinetics and maturation stage from a Vδ2 cell number >30 cells.

**Table 1.** Patient characteristics.

| Clinical status | Patients |  |  | Healthy volunteers |  |
| --- | --- | --- | --- | --- | --- |
|  | First wave | Second wave | Post COVID | First wave | Second wave |
| Number of patients | 6 | 4 | 7 | 7 | 4 |
| Median age, years | 64 (55-74) | 60 (56-63) | 57 (45-60) | 34 (30-30) | 30.5 (26-36) |
| Gender (M/F) | 4/2 | 3/1 | 3/4 | 2/5 | 1/3 |
| Overweight | 2 (33) | 1 (25) | 4 (57) | x | x |
| Obesity, n (%)* | 0 (0) | 1 (25) | 2 (29) | x | x |
| Diabetes, n (%) | 2 (33) | 2 (50) | 2 (29) | x | x |
| Hypertension, n (%) | 2 (33) | 0 | 2 (29) | x | x |
| Hospitalization, n (%) | 6 (100) | 4 (100) | 4 (57) | 0 (0) | 0 (0) |
| Median time from symptom onset to hospital admission, days | 6 (3-7) | 6 (4-8) | 13 (10-16) | - | - |
| Median time from symptom onset to first sample, days | 7 (5-9) | 6 (5-9) | 84 (78-104) | - | - |

| Maximal WHO Clinical<br>Progression Scale** |  |  |  |  |  |
| --- | --- | --- | --- | --- | --- |
| WHO score 1-4, n (%) | 4 (67) | 0 | 3 (43) | - | - |
| WHO score 5, n (%) | 1 (17) | 4 (100) | 3 (43) | - | - |
| WHO score 6, n (%) | 1 (17) | 0 | 1 (14) | - | - |
| Median length of hospital stay<br>(range), days | 14 (9-17) | 4 (4-5) | 9 (7-10) | - | - |
| Steroids, n (%) | 1 (17) | 3 (75) | 3 (43) | 0 (0) | 0 (0) |
| Hydroxychloroquine, n (%) | 0 (0) | 0 | 4 (57) | 0 (0) | 0 (0) |

\* Obesity is defined as BMI >30 kg/m<sup>2</sup>
\*\* Refers to the maximal respiratory support required during the hospital/home stay with score 1-4 indicating no oxygen; score 5 indicating oxygen by nasal prong or facial mask; and score 6 indicating high flow oxygen therapy or non-invasive ventilation.

For each specific cell type, the panel included the differentiation markers CD27 and CD45RA to identify four maturation subsets: naive (Tn, CD27^+^ CD45RA^+^), central-memory (Tcm, CD27^+^ CD45RA^−^), effector-memory (Tem, CD27^−^ CD45RA^−^) and terminal effector-memory (Temra, CD27^−^ CD45RA^+^) for CD4 and CD8 T conv (**Supplemental Figure 2B and Figure 2C**) and Vδ2 T cells (**Figure 1C**). Vδ2 T cells maturation highlight a preferential accumulation of Temra population probably induced by SARS-CoV-2 (**Figure 1C**). Indeed, Temra Vδ2^+^ T cells increased from early to late infection (mean ± SD = 24.6 ± 14.8 *vs*. 32 ± 18.5, respectively), which exceeded the frequency in HV although not significantly (mean ± SD = 16.4 ± 9.7, *p*=0.4906). To further characterize Temra Vδ2 T cell subsets, we used an unsupervised hierarchical clustering with MeV (**Figure 1D**). Heatmap suggest two patient’s clusters: D0, PC, HV and D7, D14, D28. Temra Vδ2 T cells at D0 exhibited characteristics of highly differentiated T cells with low proliferative potential (CD57^+^) and strong cytotoxic function (CD56^+^ CD8^+^CD16^+^), as compared to HV. D7, D14, D28 and PC showed a less activated phenotype than D0.

### BTN3A expression and Vγ9Vδ2 T cells infiltration in the lung

Immunofluorescence staining of BTN3A or Vδ2 (**Figure 2**) was performed in normal human lung tissue (HLT), healthy tonsil (HT), and post-COVID-19 lung tissue (PILT). HT is known to contain large numbers of Vδ2 T cells and to express BTN3A, and was used as an internal positive control. BTN3A expression was increased in PILT as compared to NLT, as demonstrated by the staining mean fluorescence intensity (MFI) and 3D surface plot (MFI, 45.5 *vs*. 14.7) (**Figure 2A, B**). Second, we observed more Vδ2 T cell infiltration in PILT, as compared to NLT (42 *vs*. 2, respectively; **Figure 2C**). Taken together, these data suggest a potential increase of BTN3A expression and Vγ9Vδ2 T cell infiltration in lung tissue after SARS-CoV-2 infection.

**Figure 2.**
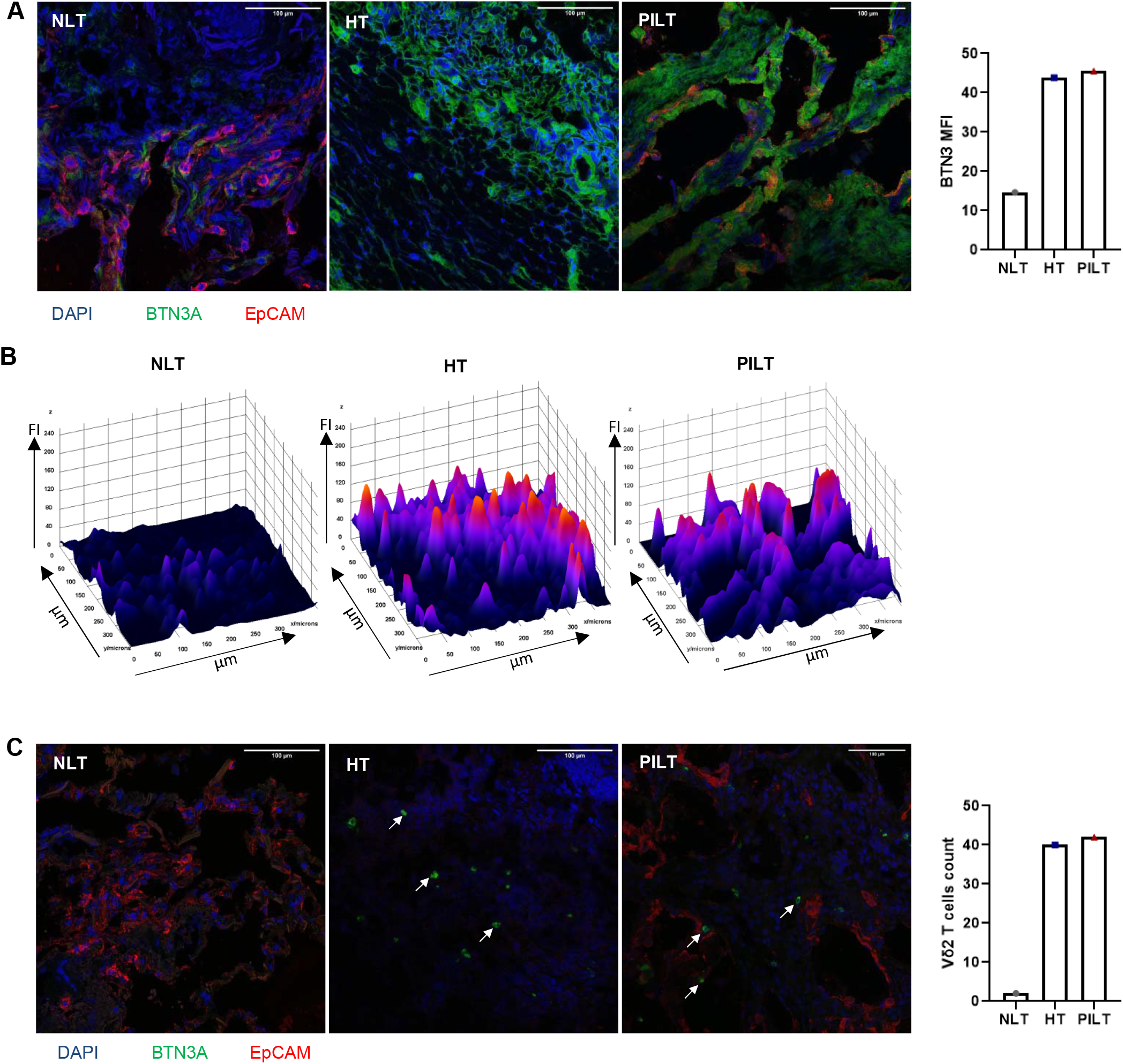
BTN3A expression and Vγ9Vδ2 T cells infiltrate in the lung. IHF analysis by confocal microscopy illustrates the spatial distribution of BTN3A (**A**), the intensity of BTN3A staining (FI = fluorescence intensity, **B**) and Vγ9Vδ2 T cells (**C**) on Normal Lung Tissue (NLT), Healthy Tonsil (HT) and Post-Infection Lung Tissue (PILT) by SARS-CoV-2 from left to right.

### Activation of Vγ9Vδ2 T cells

The kinetics of 4 patients with acute infection included during the second wave of the COVID-19 pandemic (second cohort) was studied between D0 and D90. Baseline characteristics of the patients are presented in **Table 1**.

We investigated the dynamic of the different immune cell subsets using multicolor spectral cytometry panel which identifies NK cells, CD4 and CD8 T cells and specifically study γδT cells population (**Supplemental Table 2**). The cellular responsiveness to BTN3A-mediated activation was tested using an anti-BTN3A monoclonal antibody (20.1) that selectively activates Vγ9Vδ2 T cells, which was compared with a positive control (PMA + ionomycin) and an unstimulated control (**Figure 3A**). The dimensionality reduction by UMAP analysis of CD4, CD8, and Vγ9Vδ2 T cells, and NK cells (**Figure 3B**) showed a distribution of the main lymphocyte’s subsets according to marker co-expressions (**Supplemental Figure 3**). Differentiation markers were used to characterize the maturation profiles of patients, which suggested that Temra Vδ2 T cells though highly differentiated (**Figure 3C**) tended to be more abundant in this cohort (58.4 ± 36.4 at baseline D0) as compared to the first cohort (24.6 ± 14.8 at baseline D0). Unsupervised hierarchical clustering with MeV (**Figure 3D**) classified two patient clusters: 1) D0, D7, D14, and 2) D28, D90. Interestingly, HV clustered with D28 and D90. We observed that Temra Vδ2 T cells in acute infection (D0, D7, and D14) have a very active profile with strong cytotoxic function (CD16^+^ CD8^+^CD56^+^), as compared to Temra Vδ2 T cells in late-stage of infection or post infection (D28, D90) and HV, which exhibit a senescent phenotype with high expression of CD57. We observed that anti-BTN3A monoclonal antibody activates Vγ9Vδ2 T cells to produce TNFα (Tcm: D0 *vs*. D14 p=0.0408, D14 *vs*. HV *p*=0.0328 and Tem: D14 *vs*. HV *p*=0.0043) and IFNγ (Tem: D14 *vs*. HV p=0.0298) pro-inflammatory cytokines. We also observed a degranulation response with CD107ab (Tcm: D14 *vs*. HV p=0.0131 and Tem: D14 *vs*. HV p=0.0027) (**Figure 4**). Thus, Temra Vδ2 T cells seem to be as functional as the other maturation profiles with peak effects at D14 (TNFα: 70 ± 4.1, IFNγ: 43.9 ± 3.9, CD107ab: 57.9 ± 11.9) and a consistently higher expression at D90 (TNFα: 64.6 ± 2.9, IFNγ: 37.7 ± 2.6, CD107ab: 49.6 ± 12.1), as compared to HV (TNFα: 46.9 ± 5, IFNγ: 23.7 ± 4.3, CD107ab: 32.5 ± 7.8) although these did not reach statistical significance. These modifications of the Vγ9Vδ2 T cells suggested that these cells might play a role in the recognition and killing of SARS-Cov-2 infected cells.

**Figure 3.**
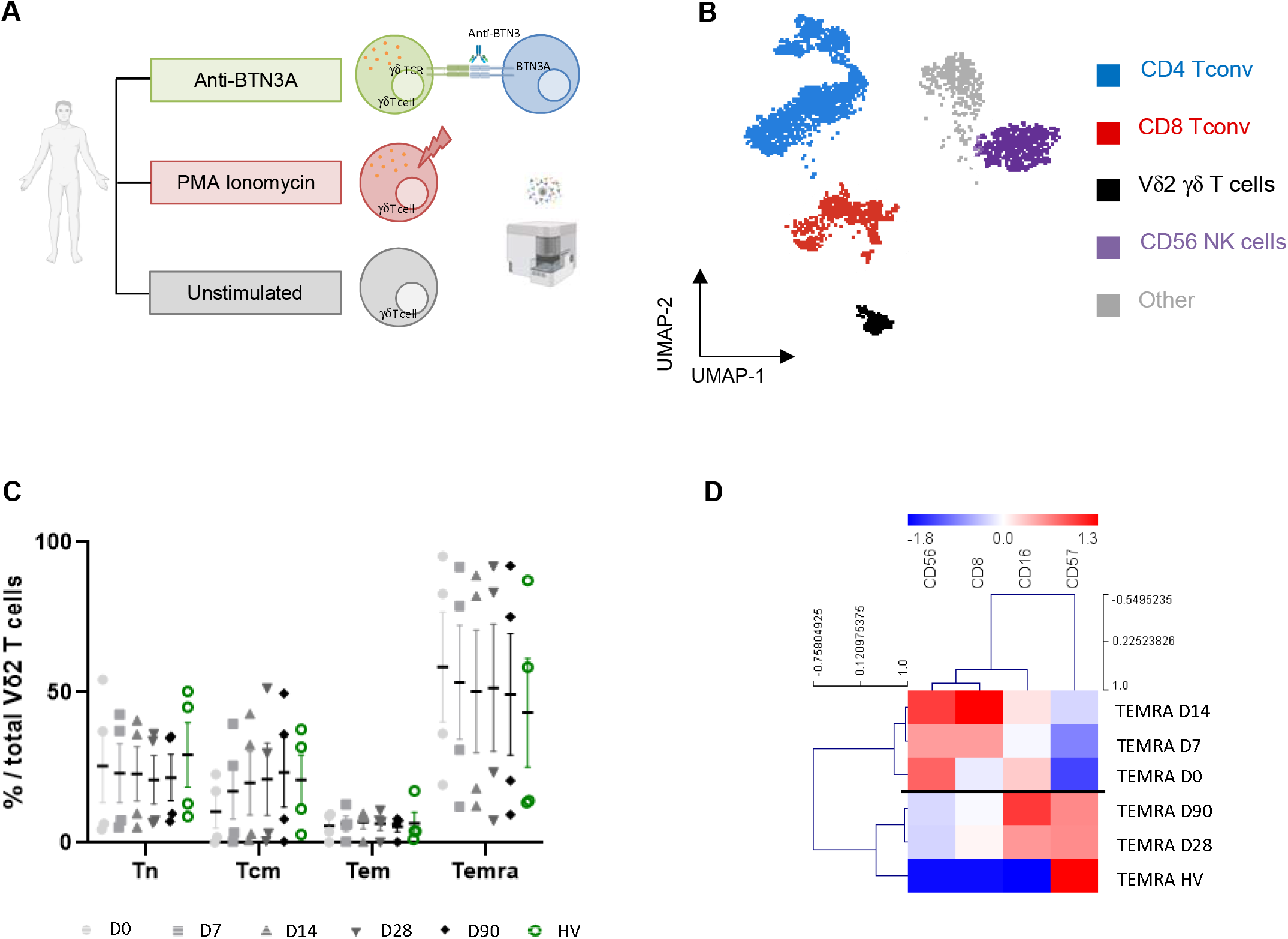
Characterization of circulating Vδ2 γδ T cells in second cohort. (**A**) Workflow. Anti-BTN3A selectively activates Vγ9Vδ2 T cell unlike PMA and ionomycin stimulation bypasses the T cell membrane receptor complex and will lead to activation of several intracellular signaling pathways. In both cases, resulting in Vγ9Vδ2 T cell activation with production of cytokines (TNFα, IFNγ) and cytotoxicity activity (CD107ab) highlighted by spectral cytometry (Cytek®Aurora). (**B**) Manual gating clusters projected on to UMAP in OMIQ software. Subsampling with 1,000 cells of D90 consensus file (n=4) was illustrated. The following markers were used to identify the immune cell populations: CD16 and CD56 for NK cells (purple), CD3, CD4 (blue) and CD8 (red) for the main T cell populations, Vδ2 TCR for Vδ2 T cells (black) and other cells (gray) which are probably B cells. (**C**) Dot plot of maturation Vδ2 γδ T cell subpopulations, identified by manual gating using CD27 and CD45RA markers. Results were showed for the patients (gray-scale kinetics) and HV (green). Data were analyzed using two-way ANOVA and Holm-Sidak’s multiple comparisons test. (**D**) Shown representative heatmap of Temra Vγ9Vδ2 T cells maturation. Blue indicates low cellular frequency and red indicates high cellular frequency. The dendrograms for markers (columns) and samples (rows) were constructed with hierarchical clustering (Pearson’s correlation). Normalized expressions of each marker are from averages by kinetics and maturation stage from a Vδ2 cell number >30 cells.

**Figure 4.**
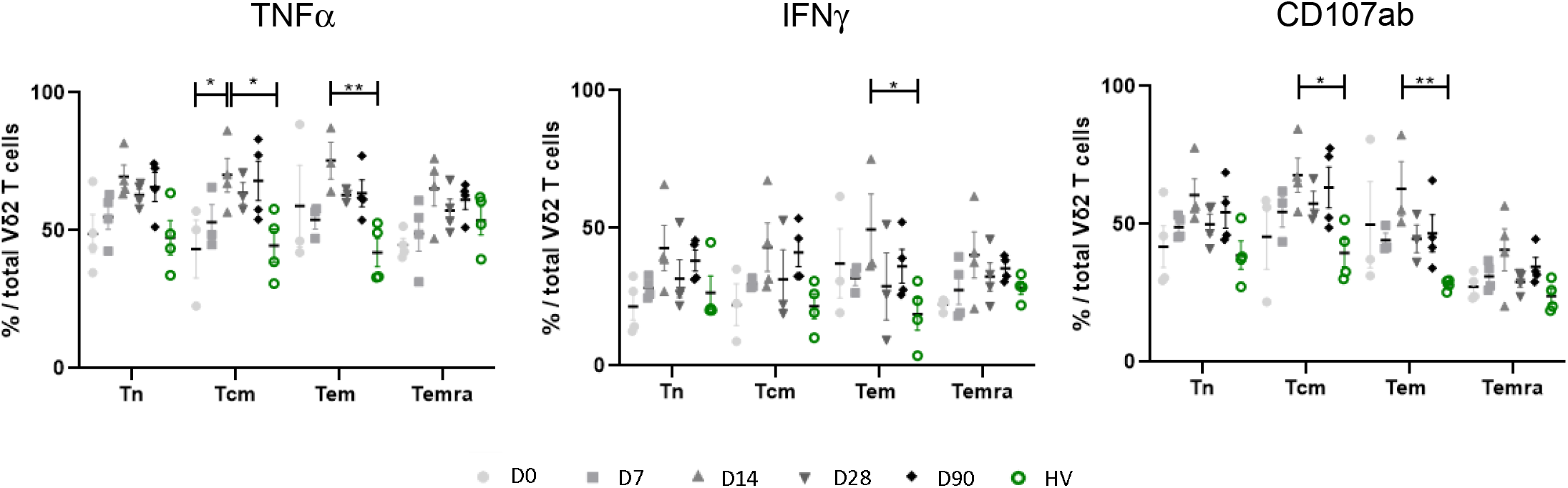
Activation of Vγ9Vδ2 T cells. Functional assay was performed in samples from patients at multiple time points (n=4) and from HV (n=4). PBMCs were stimulated with BTN3A 20.1 for 4 hours. Response was assessed by measuring cells positive for TNFα, IFNγ and CD107ab according to the maturation profile of Vγ9Vδ2 T cells. Data were analyzed using two-way ANOVA and Holm-Sidak’s multiple comparisons test \**p*<0.05 and \*\**p*<0.01.

### Effects of SARS-CoV-2 infection on BTN3A expression and characterization of Vγ9Vδ2 T cells responses against SARS-CoV-2 infected cells in vitro

First, we evaluated whether SARS-CoV-2 infection affected the expression of BTN3A on cells from myeloid primary cell cultures of monocytes or macrophage derived from monocytes (MDMs) or non-tumorigenic lung cell lines (BEAS-2B or MRC-5). Gene expression of the three isoforms (A1, A2, A3) of *BTN3A* was significantly increased 5-fold in SARS-CoV-2 stimulated MDMs, as compared to unstimulated MDMs (A1 *p*=0.012, A2 *p*=0.006, A3 *p*=0.028) (**Figure 5A**). No statistically significant difference in *BTN3A* transcriptional levels was observed in monocytes or lung cells cultures. However, BTN3A protein expression was increased 3.5-fold in MDMs and 2-fold in BEAS-2B and MRC-5 cells following stimulation with SARS-CoV-2, as compared to controls (*p*=0.048, *p*=0.012 and *p*=0.002, respectively) (**Figure 5B**).

**Figure 5.**
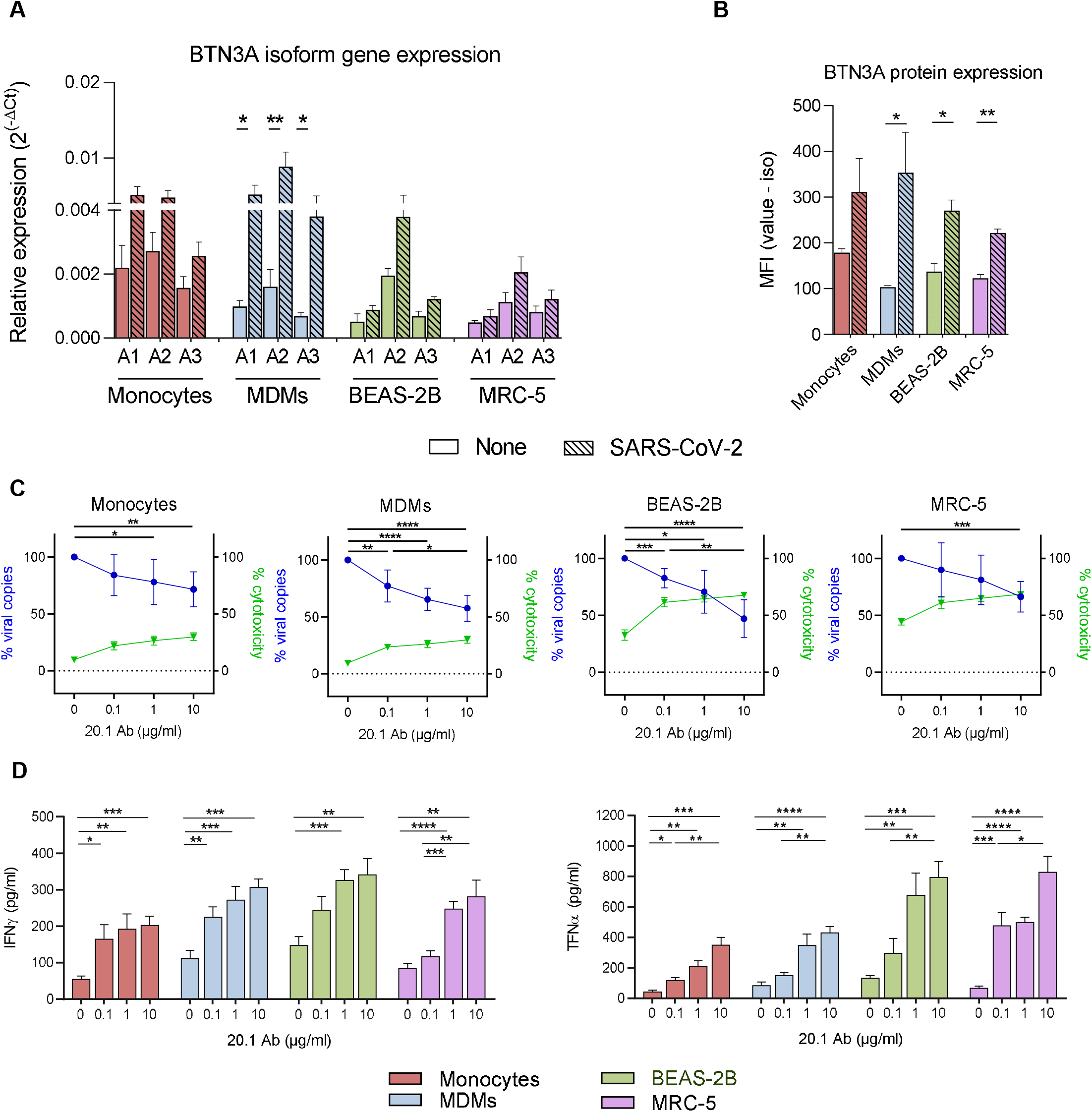
BTN3A expression in response to SARS-CoV-2 infection in monocytes, MDMs and two lung non-tumorigenic cell lines and assessment of Vγ9Vδ2 T lymphocytes anti-SARS-CoV-2 responses. (**A, B**) Monocytes, MDMs, BEAS-2B and MRC-5 cells were stimulated with SARS-CoV-2 IHU-MI6 strain (1 MOI) for 24 hours. (**A**) Mean fluorescence intensity (MFI) of BTN3A expression was investigated for monocytes, MDMs, BEAS-2B and MRC-5 cells (n=3). (**B**) The gene expression of the three *BTN3A* isoforms (A1, A2, A3) was investigated by qRT-PCR after normalization with housekeeping gene as endogenous control. Relative quantity of investigated genes at 24 hours of stimulation was evaluated for monocytes and MDMs (n=6), and BEAS-2B and MRC-5 cells (n=3). (**C, D**) Monocytes, MDMs, BEAS-2B and MRC-5 cells (n=6) were stimulated with SARS-CoV-2 IHU-MI6 strain (1 MOI) and co-cultured with Vγ9Vδ2 T cells at effector-to-target (E:T) ratio of 1:1 for 24 hours in presence of anti-BTN3A 20.1 Ab (0, 0.1, 1 or 10 μg/ml). (**C**) After 24 hours, SARS-CoV-2 viral load was quantitated by RT-PCR. Data were analyzed by the following formula 2^^–(Ct anti-BTN3A ICT01 – Ct diluent^ and represented in % viral copies. The cytotoxicity was assessed by flow cytometry as the percentage of Caspase 3/7^+^ cells in the target cell population. (**D**) The culture supernatants from co-cultures were analyzed for the presence of TNFα and IFNγ by ELISA. Values represent mean ± standard error. \**p*<0.05, \*\**p*<0.01, \*\*\**p*<0.001 and \*\*\*\**p*<0.0001.

After confirming that the viral infection did not affect Vγ9Vδ2 T cells viability (**Supplemental Figure 5**), we co-cultured Vγ9Vδ2 T cells with myeloid or lung cells in presence of SARS-CoV-2 IHU-MI6 strain and increasing concentrations of 20.1 mAb, which produced a dose-dependent inhibition of viral replication (28.4% of viral replication inhibition in monocytes cultures *p*=0.0011, 42.4% in MDMs *p*<0.0001, 53.0% in BEAS-2B *p*<0.0001 and 33.7% in MRC-5 cultures *p*=0.0001 at 10 μg/ml) (**Figure 5C**). Cytotoxicity of Vγ9Vδ2 T lymphocytes appeared to be higher towards SARS-CoV-2 infected lung cells than towards myeloid cells (9.6% in monocytes, 9.3% in MDMs, 32.7% in BEAS-2B and 44.4% in MRC-5). The anti-SARS-CoV-2 cytotoxicity of Vγ9Vδ2 T cells increased along with concentrations of anti-BTN3A antibody (+20.2% of cytotoxicity in monocytes cultures, +20.5% in MDMs, +35.0% in BEAS-2B and +24.1% in MRC-5 at 10μg/ml, *p*<0.0001 for each cell cultures).

Finally, we investigated if Vγ9Vδ2 T cells activated with 20.1 mAb exerted a non-cytolytic anti-SARS-CoV-2 activity based on cytokine production. Increasing concentrations of anti-BTN3A were associated with a significant and dose-dependent increase of both IFNγ and TNFα in the supernatants of the four cell cultures (difference in mean ± SEM between 0 and 10 μg/ml of 20.1 mAb for IFNγ: monocytes 147.2 ± 27.5 *p*=0.0003, MDMs 194.3 ± 32.5 *p*=0.0001, BEAS-2B 194.2 ± 51.9 *p*=0.0038 and MRC-5 196.9 ± 47.9 *p*=0.0021 and for TNFα: monocytes 307.4 ±54.8 *p*=0.0002; MDMs 346.7 ± 47.6 *p*<0.0001, BEAS-2B 661.7 ± 108.3 *p*=0.0001 and MRC-5 761.8 ± 105.5 *p*<0.0001) (**Figure 5D**).

## DISCUSSION

Anti-SARS-CoV-2 immune responses and more specifically the involvement of innate immunity in the disease course, severity, symptoms chronicity and recovery are far from being completely elucidated. Improving our knowledge on COVID-19 immune responses is needed to allow a better management of the disease and drive the development of novel therapeutic approaches. Vγ9Vδ2 T cells are important effectors of innate immunity. They constitute the predominant subset among circulating γδ T cells and are known to sense stress-signals in infected or transformed cells linking innate to adaptive responses^15^. Their involvement in COVID-19 immune responses have been scarcely described in previous publications^8,9^. In this study, we sought to further understand the features of Vγ9Vδ2 T cells responses against the SARS-CoV-2 infection.

Our phenotyping results on peripheral blood from patients with SARS-CoV-2 infection, strikingly demonstrates a dramatic decrease in Vγ9Vδ2 T lymphocytes. This profound decrease also observed in case of mild infection seems to persist for a prolonged period in PC patients. This observation differs from the *Mycobacterium tuberculosis* infections both in humans and cattle that are associated with an expansion of Vγ9Vδ2 T lymphocytes^11,20^. Our hypothesis is that Vδ2 T cells might be recruited toward the infected tissues like the lung as already shown in humans after *M. tuberculosis* infections^20^. Thus, we have showed with lung samples a potential Vγ9Vδ2 T cells trafficking in PILT. Furthermore, we also observed overexpression of BTN3A in PILT a key activator of Vγ9Vδ2 T cells suggesting their potential implication to kill SARS-CoV-2 infected cells. Unlike the first cohort, the second cohort did not presented this shift in Vδ2 T cells frequency; different clinical characteristics between the two cohorts including the more common use of steroids or another SARS-CoV-2 variant could potentially explain this difference. Moreover, a striking feature was a rapid, meaning as soon as the patients entered the hospital, differentiation towards the terminally differentiated Temra (CD45RA^+^ CD27^−^) subset meaning the one detected in the blood and tissues. Previous studies have already highlighted the role of Vγ9Vδ2 T cells during SARS-CoV-2. Indeed, Rijkers et al. described a reduction Vγ9Vδ2 T cell numbers in SARS-CoV-2 infected patients with fatal outcome. As seen with SARS-CoV-1^13^, the Vγ9Vδ2 T cell population in surviving patients shifted to effector memory phenotype^8,13^. By contrast, another study has shown an increase of naïve γδ T cells, and a decrease of effector γδ T cells after mild and severe COVID-19 infected patients upon entry at the hospital^9^. In addition to these results, Temra Vδ2 T cell subset presented high activated phenotype (CD16^+^ CD8^+^ CD56^+^) especially for the second cohort confirmed by functional assay with the anti-BTN3A activating mAb. Temra Vδ2 T cells have both cytotoxic and pro-inflammatory activities through CD107ab and IFNγ, TNFα expression, a phenomenon that is likely to contribute to the effective anti-viral response.

Our study was limited by the assessment of a limited cohort, thus data deserves to be completed for a better understanding of the mechanistic of Vγ9Vδ2 T cells immune responses during and after COVID-19 recovery in severe and mild disease and in chronic COVID-19 patients. However, the ability to follow patients weekly increases the liability of these data. It will be also important to investigate the evolution of Temra population following re-infection and the current vaccine campaigns.

In the second part, one of the main unprecedented findings of our study is the up-regulation by the SARS-CoV-2 infection of the expression of BTNA3, a plasma membrane protein required to initiate Vγ9Vδ2 T cells activation, on myeloid and lung cells. Macrophages are the cells whose BTN3A expression is most strongly modulated by infection. Indeed, we observe a significant increase in the surface expression but also in the transcriptomic expression of the 3 isoforms *BTN3A1, BTN3A2* and *BTN3A3*. The fact that the molecule is more expressed following infection could enhance the Vγ9Vδ2 T cell activation whose viability is not affected by infection and thus their antiviral activity. Therefore, we used the anti-BTN3A activating mAb in the Vγ9Vδ2 T-cell/ infected cell co-cultures to evaluate its impact on the antiviral activity against SARS-CoV-2. We demonstrated that Vγ9Vδ2 T cells can elicit *in vitro* strong cytotoxic and non-cytolytic IFN-γ dependent anti-SARS-CoV-2 activities when activated with an anti-BTN3A antibody which is consistent with findings from previous studies reporting on antiviral activities of γδ T cells^21,22^.

The COVID-19 pandemic has led to substantial mortality worldwide. Despite promising vaccination campaigns, the potential emergence of new strains which might limit vaccines effectiveness remains a major threat. Therefore, the need to develop anti-SARS-CoV-2 therapies is urgent. Immunotherapies to overcome immune responses imbalances in some patients will be needed together with other antiviral drugs to implement precision medicine approaches. Immunotherapies triggering Vγ9Vδ2 T cells responses either through the direct use of amino-biphosphonates or through the adoptive transfer of ex-vivo activated Vγ9Vδ2 T cells have been discussed in literature^21,23^. The use of anti-BTN3A monoclonal antibodies to elicit Vγ9Vδ2 T cells activities, such as ICT01 that is being currently investigated in cancer patients (NCT04243499, ImCheck Therapeutics), could be a promising therapeutic option.

In summary, we observed a prolonged and dramatic decrease in Vγ9Vδ2 T lymphocytes on peripheral blood from patients with SARS-CoV-2 infection. Most of the Vγ9Vδ2 T cells were terminally differentiated Temra (CD45RA^+^ CD27^−^) subset which exhibited high activated phenotype (CD16^+^ CD8^+^ CD56^+^). Finally, we discovered that the BTN3A molecule was modulated on myeloid cells and lung cell lines following SARS-CoV-2 infection. Interestingly, the use of anti-BTN3A activating antibody, that enables the activation of Vγ9Vδ2 T cells leads to the inhibition of SARS-CoV-2 replication in co-cultures of infected cells with Vγ9Vδ2 T cells. Whether this pathway is important to be involved in patients’ care needs to be investigated.

## ACKNOWLEDGMENTS

Laetitia Gay was supported with a Cifre fellowship with ImCheck Therapeutics. This work was supported by the IMMUNO-COVID project managed by the “Agence Nationale de la Recherche” Flash Covid (reference: IMMUNO-COVID).

This work was performed with the support of Cytek Biosciences which provided panel design and the necessary antibodies for the spectral cytometry part. Thanks to Olivier Jean for its application support and advices in spectral cytometry along the project. Thanks to Yacine Kharraz for helping with unmix reviewing. Thanks to the CRCM cytometry core facility for assistance and the Canceropôle Provence-Alpes-Côte d’Azur for its contribution to the financial support of the Aurora (Cytek Biosciences).

## AUTHORSHIP CONTRIBUTIONS

L.G, M.S.R, M.R and L.G performed the experiments and analyzed the data. S.M, E.F, L.M, P.F, C.C, J.L.M and D.O supervised the work. L.G, M.S.R, S.M, J.L.M and D.O participated in the writing of the paper. All the authors read and approved the final manuscript.

