## Supplemental data for "Vγ9Vδ2 T cells are potent inhibitors of SARS-CoV-2 replication and exert effector phenotypes in COVID-19 patients"

### TABLES

**Supplemental Table 1. Reagents used for spectral cytometry for the first cohort (phenotype)**

| Antibodies | Fluorochromes | Clones | Supplier |
|---|---|---|---|
| CD45RA | BUV395 | 5H9 | BD Biosciences |
| TCR Vδ2 | BUV615 | B6 | BD Biosciences |
| CD56 | BUV737 | NCAM16-2 | BD Biosciences |
| CD8 | BUV805 | SK1 | BD Biosciences |
| CCR7 | BV421 | G043H7 | Biolegend |
| CD3 | BV510 | OKT3 | Biolegend |
| CD20 | Pacific Orange | HI47 | Life technologies |
| CD33 | BV570 | WM53 | Biolegend |
| CD28 | BV650 | CD28-2 | Biolegend |
| CD57 | FITC | HNK-1 | Biolegend |
| CD14 | Spark Blue 550 | 63D3 | Biolegend |
| CD45 | PerCP | 2D1 | Biolegend |
| TCRpanγδ | PerCP eFluor710 | B1-1 | Invitrogen |
| CD4 | CF568 | SK3 | Cytek |
| CD25 | PE | BC96 | Biolegend |
| CD27 | APC | M-T271 | Biolegend |
| CD127 | APC-R700 | HIL-7R-M21 | BD |
| CD19 | Spark NIR 685 | HIB19 | Biolegend |

**Supplemental Table 2. Reagents used for spectral cytometry for the second cohort (function)**

| Antibodies | Fluorochromes | Clones | Supplier |
|---|---|---|---|
| CD45RA | BUV395 | 5H9 | BD Biosciences |
| CD16 | BUV496 | 3G8 | BD Biosciences |
| TCR Vδ2 | BUV615 | B6 | BD Biosciences |
| CD56 | BUV737 | NCAM16-2 | BD Biosciences |
| CD8 | BUV805 | SK1 | BD Biosciences |
| CCR7 | BV421 | G043H7 | Biolegend |
| CD57 | BV510 | QA17A04 | Biolegend |
| CD28 | BV650 | CD28-2 | Biolegend |
| IFNγ | FITC | 45.15 | Beckman Coulter |
| Vδ1 | PerCP-Vio700 | REA173 | Miltenyi |
| CD45 | PerCP | 2D1 | Biolegend |
| CD4 | CF568 | SK3 | Cytek |
| CD3 | ECD | UCHT1 | Beckman Coulter |
| TCRpanγδ | PE-Cy5.5 | IMMU510 | Beckman Coulter |
| TNFα | APC | REA656 | Miltenyi |
| CD27 | APC-Cy7 | O323 | Biolegend |
| CD127 | APC-R700 | HIL-7R-M21 | BD Biosciences |

**Supplemental Table 3. Primers**

| Gene | Forward primer (5'-3') | Reverse primer (5'-3') |
|---|---|---|
| *ACTB* | GGAAATCGTGCGTGACATTA | AGGAGGAAGGCTGGAAGAG |
| *BTN3A1* | TTCCAGGTCATAGTGTCTGC | TGAGCAGCTGAGCAAAAGG |
| *BTN3A2* | TGGGAATACCAAGGGA | AGTGAGCAGCTGGACCAAGA |
| *BTN3A3* | GAGGGAATACTAAGAAATGGT | GAAGAGGGAGACATGAAAGT |

### FIGURE LEGENDS

**Supplemental Figure 1. UMAP and manual gating to identify Vγ9Vδ2 T cells for the first cohort (phenotype)**

(**A**) UMAP to visualize each marker expression. (**B**) Manual gating to identify Vγ9Vδ2 T cells. Manual gating to identify Vγ9Vδ2 T cells. Live immune cells were pre-gated as live dead blue$^{-}$ CD45$^{+}$ and CD33$^{+}$ myeloid cells and CD20$^{+}$ CD19$^{+}$ B cells were excluded. CD3$^{+}$ TCRVδ2$^{+}$ was then identified.

**Supplemental Figure 2. Frequency of population control and maturation of T conv**

(**A**) Frequency of CD8$^{+}$ Tconv and total γδ T cells in the first cohort. (**B, C**) Maturation of T CD4 and T CD8 conv in the first cohort (**B**) and the second cohort (**C**). We observed one patient with increased Temra CD4 Tconv frequency, this was a patient with probably another infection.

**Supplemental Figure 3. UMAP and manual gating to identify Vγ9Vδ2 T cells for the second cohort (function)**

(**A**) UMAP to visualize each marker expression. (**B**) Manual gating to identify Vγ9Vδ2 T cells. Live immune cells were pre-gated as live dead blue$^{-}$ CD45$^{+}$ and myeloid cells were excluded by SSC-A vs. FSC-A. CD3$^{+}$ TCRVδ2$^{+}$ was then identified.

**Supplemental Figure 4. Microscopy control of BTN3 and Vδ2 staining**

Microscopy unstained control of BTN3 (**A**) and Vδ2 (**B**). BTN3 and Vδ2 staining (left and right, C) in PILT.

**Supplemental Figure 5. Impact of SARS-CoV-2 on Vγ9Vδ2 T cell viability**

Vγ9Vδ2 T cells (from 3 healthy volunteers) were stimulated with SARS-CoV-2 IHU-MI6 strain (0.25, 0.5 or 1 MOI). After 24 hours, the viability of Vγ9Vδ2 T cells was evaluated by flow cytometry as the percentage of live cells in the Vγ9Vδ2 T cell population. Values represent mean ± standard error of the mean.

**Supplemental Figure 1**

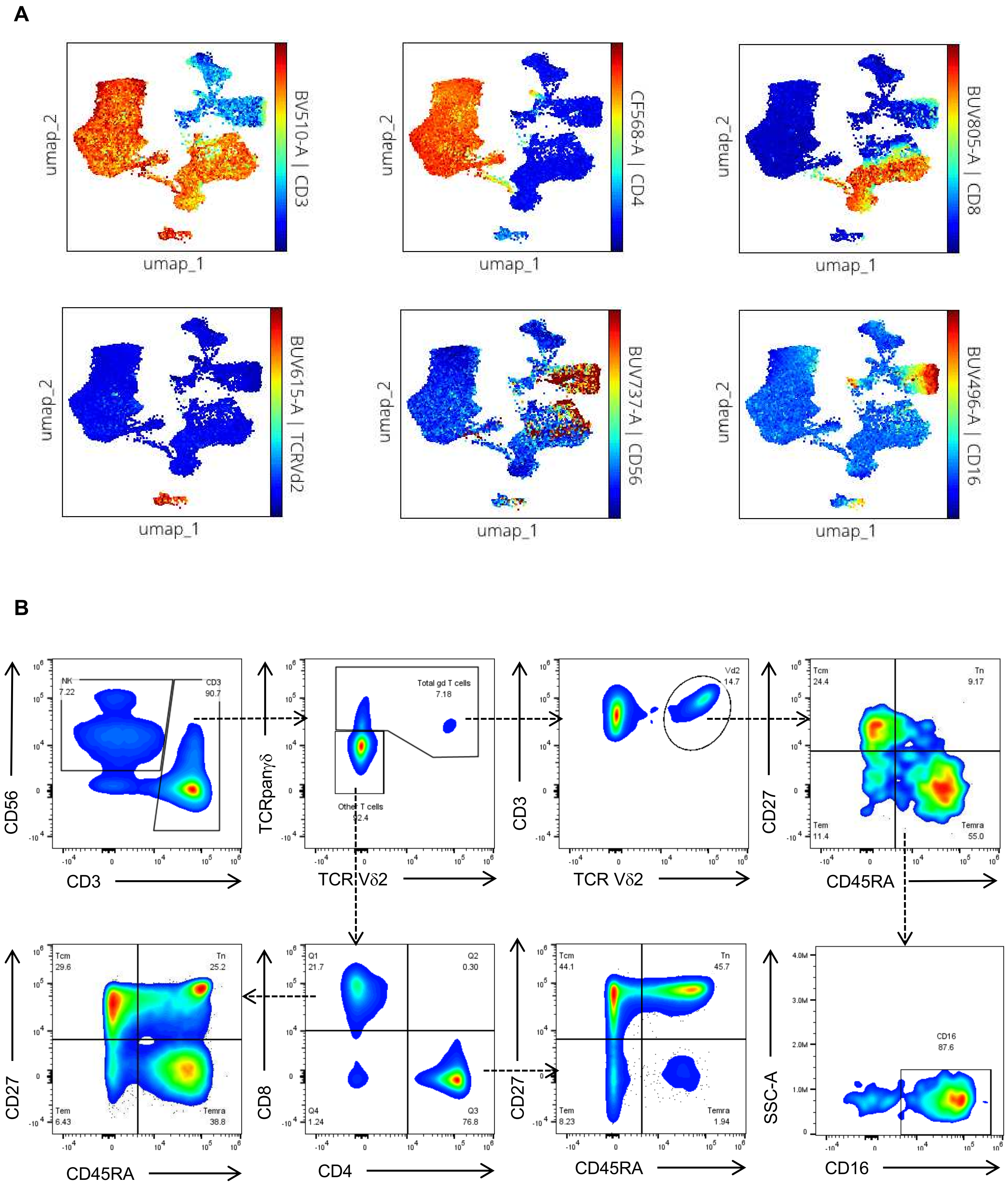

**Supplemental Figure 2**

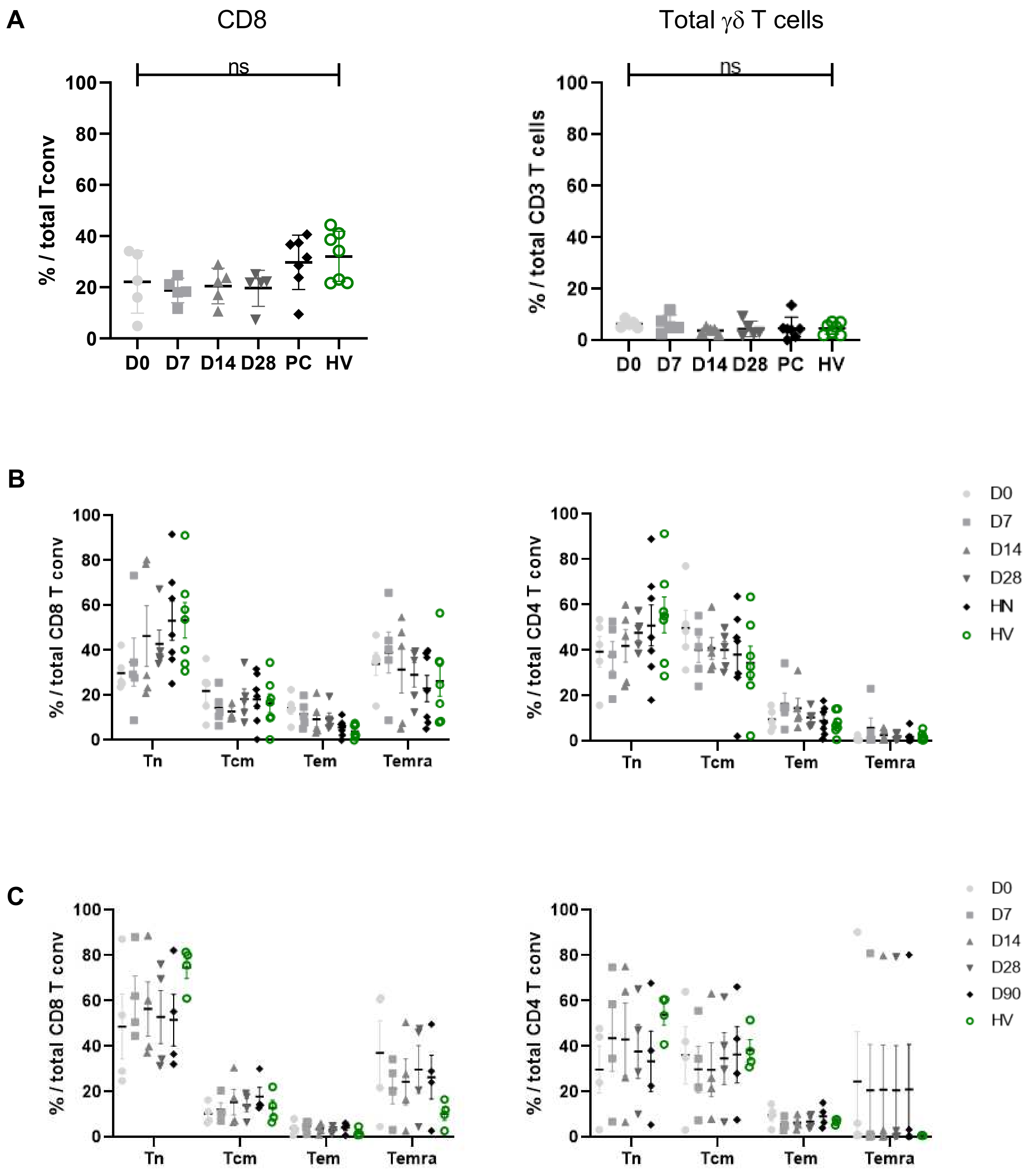

**Supplemental Figure 3**

**A**

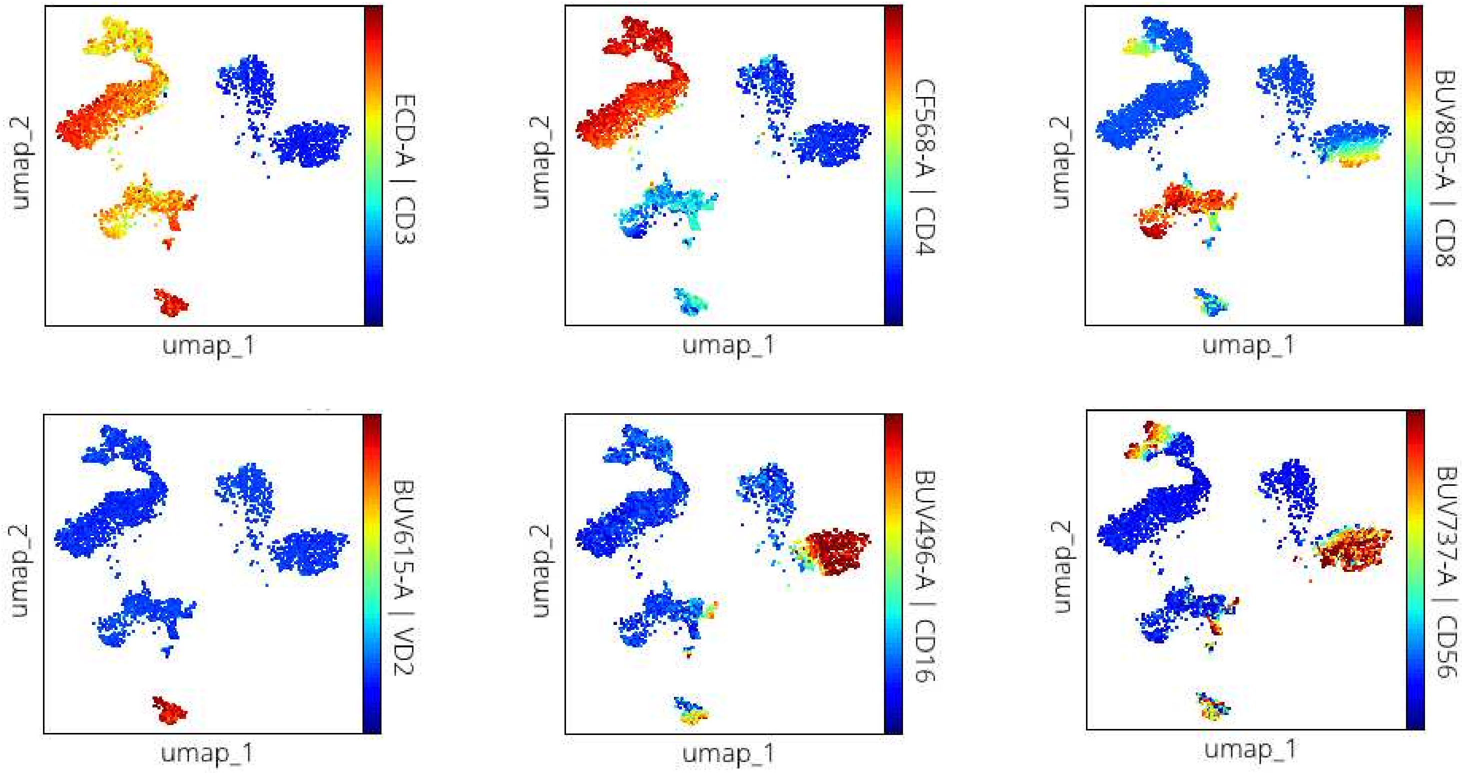


**B**

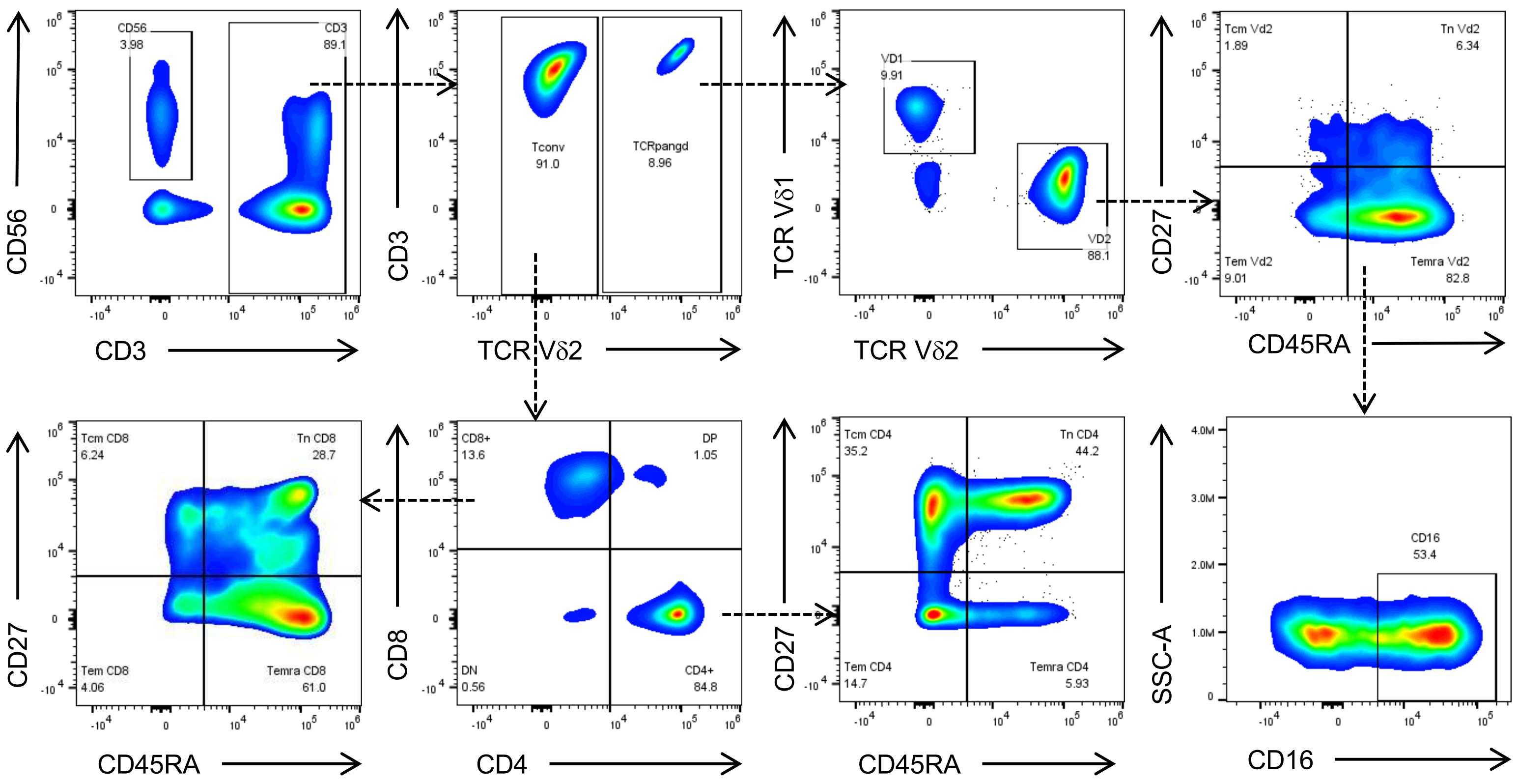

### Supplemental Figure 4

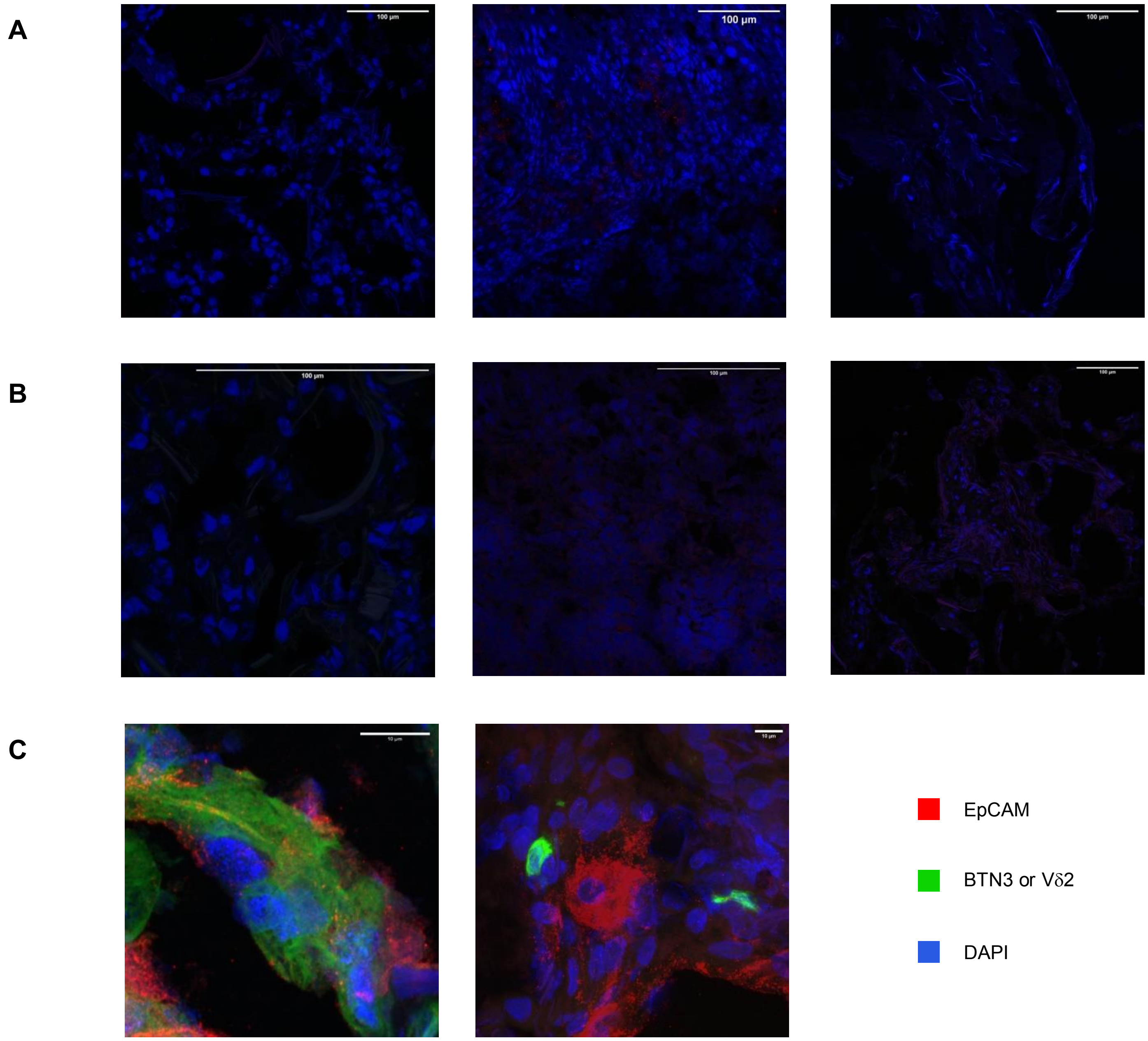

**Supplemental Figure 5**

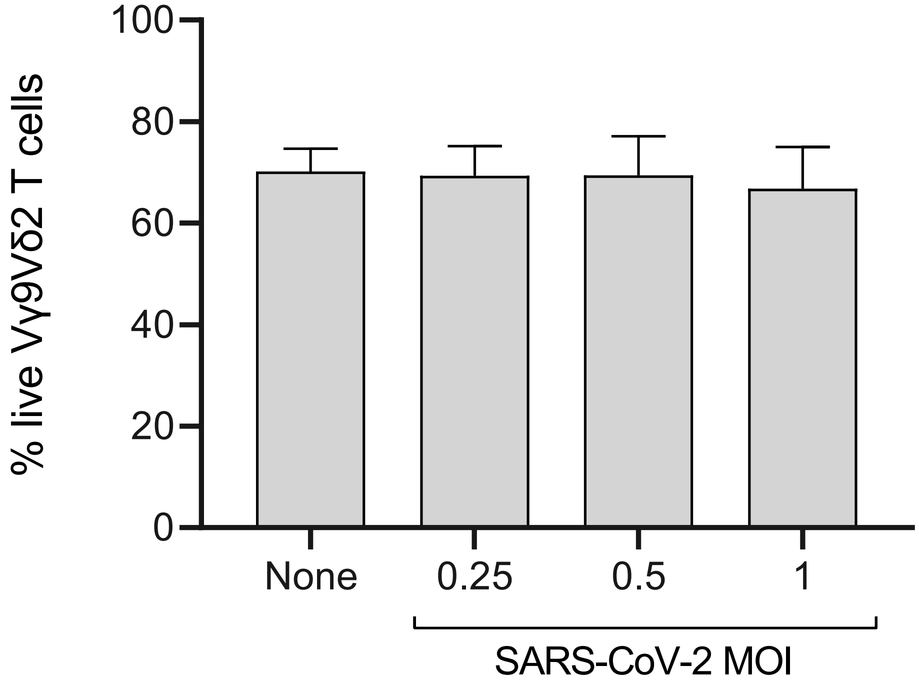
